# Profound perturbation of the human metabolome by obesity

**DOI:** 10.1101/298224

**Authors:** Elizabeth T. Cirulli, Lining Guo, Christine Leon Swisher, Naisha Shah, Lei Huang, Lori A. Napier, Ewen F. Kirkness, Tim D. Spector, C. Thomas Caskey, Bernard Thorens, J. Craig Venter, Amalio Telenti

## Abstract

Obesity is a heterogeneous phenotype that is crudely measured by body mass index (BMI). More precise phenotyping and categorization of risk in large numbers of people with obesity is needed to advance clinical care and drug development. Here, we used non-targeted metabolome analysis and whole genome sequencing to identify metabolic and genetic signatures of obesity. We collected anthropomorphic and metabolic measurements at three timepoints over a median of 13 years in 1,969 adult twins of European ancestry and at a single timepoint in 427 unrelated volunteers. We observe that obesity results in a profound perturbation of the metabolome; nearly a third of the assayed metabolites are associated with changes in BMI. A metabolome signature identifies the healthy obese and also identifies lean individuals with abnormal metabolomes – these groups differ in health outcomes and underlying genetic risk. Because metabolome profiling identifies clinically meaningful heterogeneity in obesity, this approach could help select patients for clinical trials.

## Introduction

Obesity is one of the most widespread problems facing our society’s health today. Excessive weight significantly increases risk for conditions like diabetes mellitus and cardiovascular disease^1,2^. Worldwide, the prevalence of obesity has nearly tripled since 1975, with 39% of the world’s adults being overweight and 13% being obese^3^. The high prevalence can partially be attributed to increasing consumption of hypercaloric foods and sedentary lifestyles^3^. Previous studies have identified metabolic signatures associated with obesity, including increased levels of branched-chain and aromatic amino acids as well as glycerol and glycerophosphocholines^4,5,6,7,8,9^. However, prior work has been limited by focusing on a relatively small numbers of metabolites, individuals, or obesity phenotypes.

The characterization of the metabolites that are associated with obesity can provide insights into the mechanisms that lead to this disease and associated consequences. Longitudinal assessment of weight gain and weight loss over time may indicate whether there are metabolomic changes that cause obesity—meaning that current metabolite levels could predict future weight changes—or whether all metabolite changes associated with obesity are a consequence of weight changes. Drawing genetics into this assessment allows the determination of whether genetic variation leads to metabolite changes that subsequently result in obesity, allowing the further delineation of the causal pathway to obesity. Finally, research in this area may identify biomarkers of obesity and of different types of obesity, for example biomarkers of so-called healthy obesity^10^.

There are recent calls to improve phenotyping in very large numbers of obese people with the goals of understanding factors that make people susceptible to (or protected from) obesity, accompanied by a better elucidation of the factors that account for variability in success of different obesity treatments^11^. Here, in an effort to understand the relationship between metabolic perturbations and the obese state, we analyzed 2,396 individuals with longitudinal measurements of body mass index (BMI), anthropomorphic data, whole body DEXA scans and metabolome, combined with baseline genetic risk. The metabolome assay covered up to 1,007 metabolites at up to three distinct time points for each individual over the course of study. We identified associations between nearly a third of the metabolome and BMI, and we show that metabolite levels can explain ~40% of the variation in BMI and can predict obesity status with ~80-90% specificity and sensitivity. The metabolome profile is a strong indicator of metabolic health compared to the polygenic risk assessment and anthropomorphic measurements of BMI.

## Results

### Profound perturbation of metabolome by obesity

#### Metabolites associated with BMI

We compared the levels of individual metabolites to the BMIs of 832, 882, and 861 unrelated individuals of European ancestry in the TwinsUK cohort^12^ at three timepoints spanning a total range of 8-18 years. We identified 284 metabolites that were significantly associated (p<5.5×10^−5^) with BMI at one or more timepoints (Table S1). We focused on metabolites that were significantly associated with BMI at all 3 timepoints and sought to replicate the associations in an independent sample of 427 unrelated individuals of European ancestry participating in the Health Nucleus cohort^13^. In total, our analyses identified 307 metabolites that were significantly associated with BMI in at least one cohort and timepoint (Table S1). We identified 83 metabolites that showed directions of effect that were consistent between the two cohorts, of which 49 were statistically significant replications (**Figure 1** and **Table 1**).

**Table 1.**
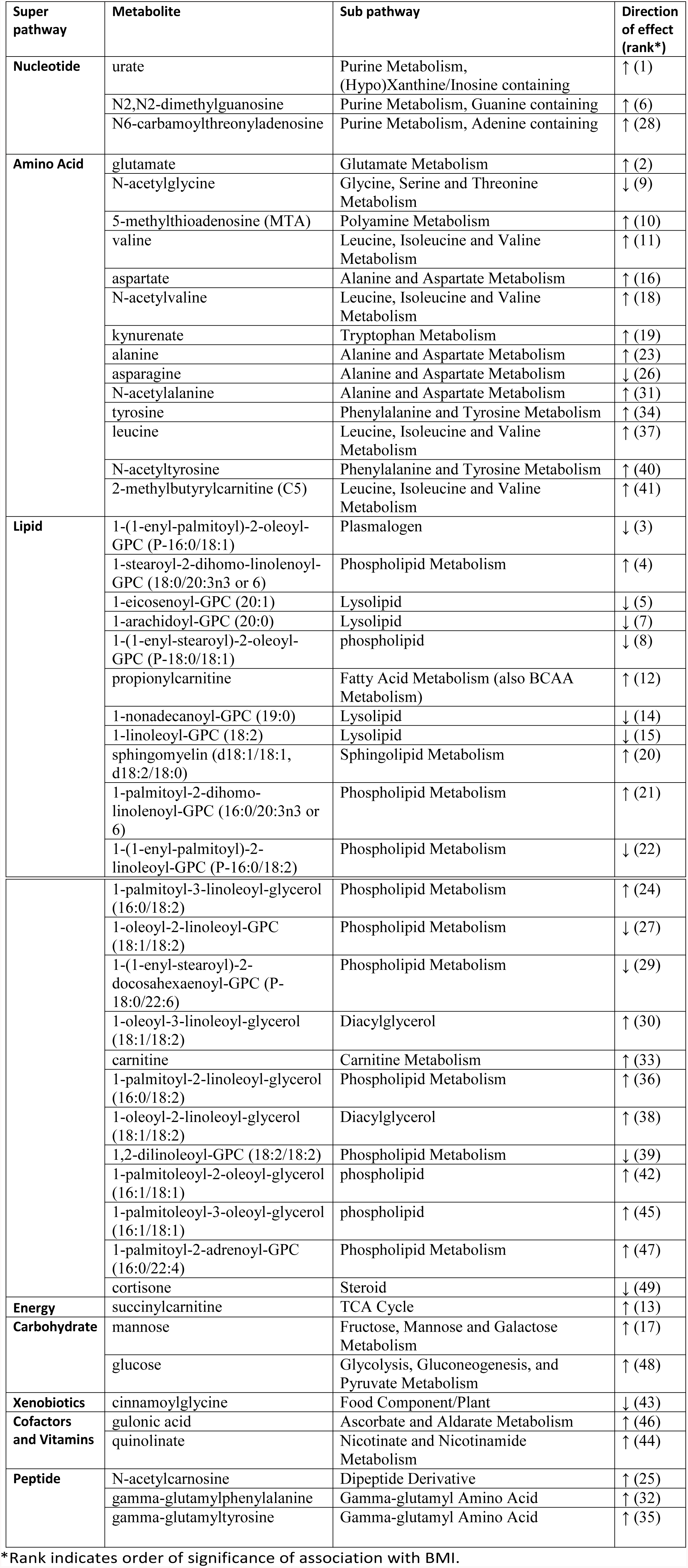
Metabolite signature associated with BMI.

**Figure 1.**
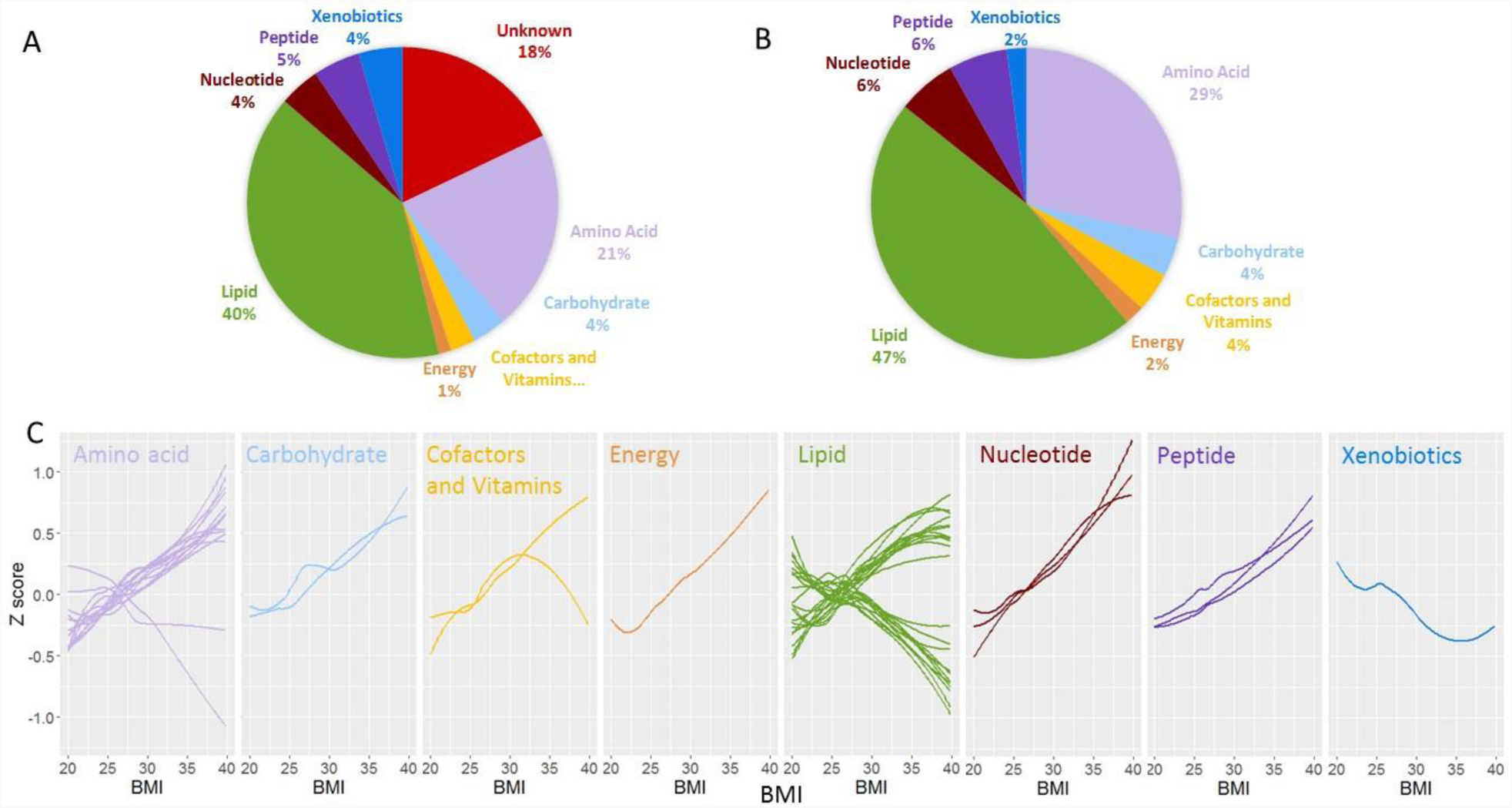
Pathway categories of metabolites associated with BMI. Shown are the pathway categories of (A) the 307 metabolites significantly associated with BMI and (B) the 49-metabolite signature. (C) The values of each of the 49 BMI-associated metabolites are plotted with a Loess curve against the BMI for timepoint 1 in TwinsUK. Only unrelated individuals of European ancestry are included, and the small number of individuals with BMI below 20 (n=31) or above 40 (n=10) are removed to keep the ends of the graphs from being skewed.

The 49 metabolites that associated with BMI were primarily lipids (n=23, accounting for 7.5% of all lipids assayed across both cohorts) and amino acids (n=14, 9.3% of all amino acids) but also included nucleotides (n=3, 12.0% of all nucleotides), peptides (n=3, 12% of all peptides), and other categories (n=6, see **Figure 1** and **Table 1**). The most significantly associated metabolite was urate (uric acid; p-value 1.2×10^−40^ for combined analysis of TwinsUK timepoint 1 and Health Nucleus data).

#### Patterns in metabolite change according to BMI

The majority of the 49 BMI-associated metabolites increased with increasing BMI (n=35) (**Figure 1**, Table S1). This included glucose, and, notably, mannose, which has recently been highlighted as playing a role in insulin resistance^14^. Most metabolites change linearly (both proportionally and inversely) with BMI, though some diverge at higher BMIs, especially 2-methylbutyrylcarnitine (see cofactors panel in **Figure 1C**). Branched-chain and aromatic amino acids as well as metabolites related to nucleotide metabolism like urate had the most rapid increases. Those that decreased (n=14) included phospholipids and lysolipids, as well as the amino acids asparagine and N-acetylglycine and the xenobiotic cinnamoylglycine, which has been identified as a product of the microbiome^15^. Of particular interest was the association with cortisone, a metabolite of the steroid hormone cortisol. We identified lower levels among the obese individuals, which is consistent with previous reports^16–19^. We examined the overall composition of the distributions of these metabolites via principal component analysis and found complex underlying correlations; in particular, the first principal component explained ~20% of the total variation in the levels of these 49 metabolites (Figure S1).

#### Modeling the metabolome of obesity

We used ridge regression to build a model that would predict BMI from the 49 BMI-associated metabolites (see **Figure 2**). We combined our data for the first visit of the TwinsUK cohort and the Health Nucleus cohort and trained with 10-fold cross-validation on a random half of the population. In our test set of the other half of the data, we found that the model could explain 39.1% of the variation in BMI (**Figure 2A**). In predicting whether participants were obese (BMI>=30) or normal weight (BMI 18.5-25), the model had an area under the curve (AUC) of 0.922, specificity of 89.1% and sensitivity of 80.2% (Figure S2). The model based on the metabolite signature was thereafter used as a tool to define mBMI, the predicted BMI on the basis of metabolome.

**Figure 2.**
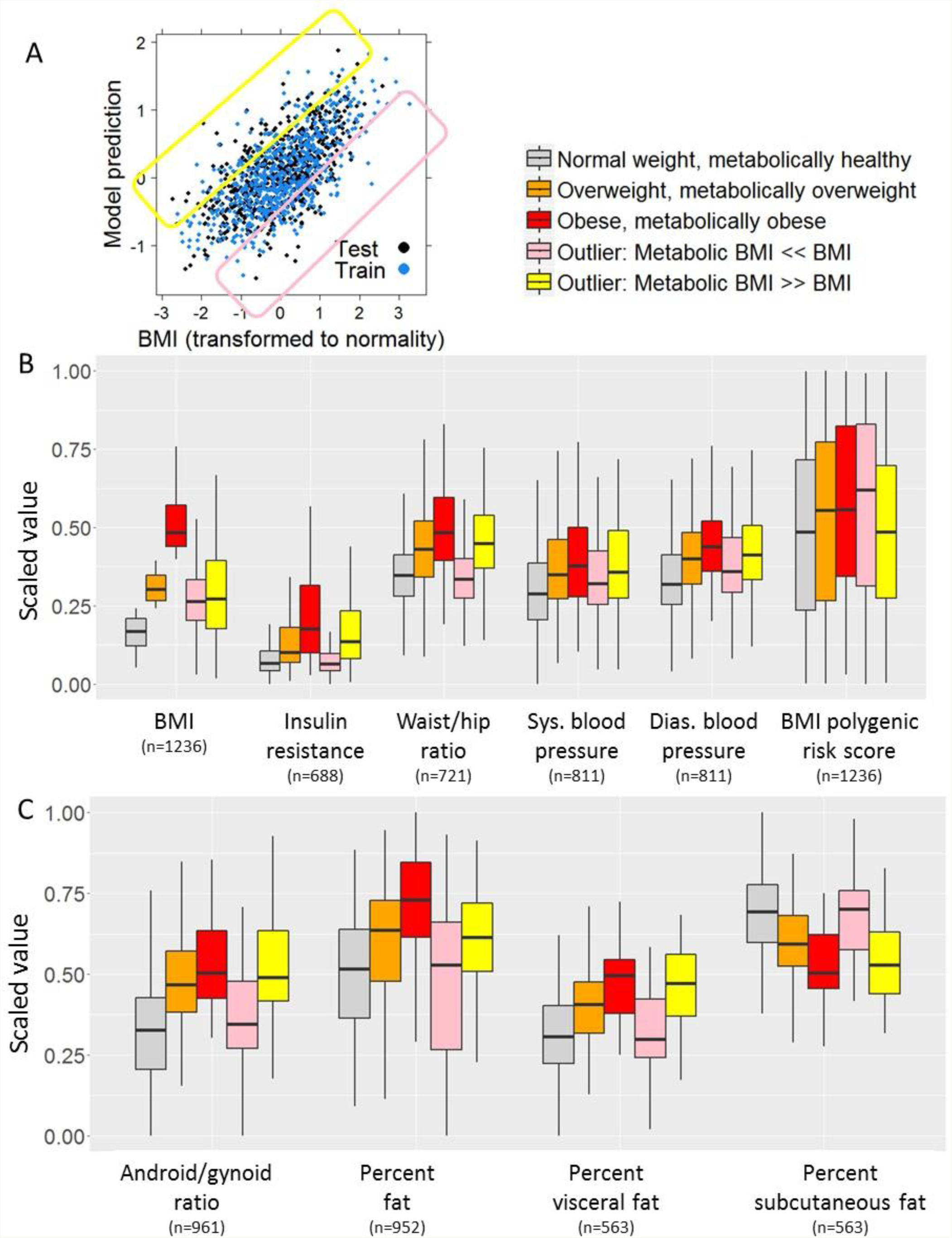
Variables associated with BMI and predicted BMI from the metabolome. (A) Correlation between ridge regression model prediction of BMI and actual BMI for all unrelated individuals of European ancestry in the TwinsUK and HN dataset. The identification of outliers is defined below: the pink box shows individuals with a much lower predicted BMI (mBMI) than actual BMI, and the yellow box shows individuals with a much higher mBMI than actual BMI. (B) Factors associated with being a mBMI outlier. Participants were split into 5 groups: those whose metabolome accurately predicted their BMI (residual after accounting for age, sex and BMI between -0.5 and 0.5) whose BMIs were either normal (18.5-25), overweight (25-30), or obese (>30); and those whose metabolome predicted a substantially higher mBMI than the actual BMI (residual <-0.5) or a substantially lower mBMI than the actual BMI (residual >0.5). All y-axis values are scaled to a range from 0-1 to allow comparison across groups. The same process is used in (C) to show DEXA imaging values associated with metabolic BMI outliers. The unexpectedly low mBMI and unexpectedly high mBMI groups had a comparable measured BMI; however, these two groups were statistically significantly different from each other (p<0.01) for all modalities except blood pressure.

### Identification and characterization of metabolic BMI outliers

Having established a model to predict BMI using the metabolome (mBMI), we split the participants into 5 groups (**Figure 2A**). Three groups included individuals whose metabolome accurately predicted their BMI (residual between -0.5 and 0.5): they were characterized as having a normal BMI (18.5-25), overweight (25-30), or obese (>30). Two groups were characterized as outliers: these included individuals whose metabolome predicted a substantially lower mBMI than the actual BMI (mBMI<<BMI, residual <-0.5) or a substantially higher mBMI than the actual BMI (mBMI>>BMI, residual >0.5). While these two outlier groups had the same weight range distribution (**Figure 2B**), they had very different values for many of the phenotypes of metabolic health collected from these cohorts (**Figure 2B&C**). Individuals with a mBMI prediction that was substantially lower than their actual BMI had levels of insulin resistance, blood pressure, waist/hip ratio, android/gynoid ratio, percent body fat, percent visceral fat, and percent subcutaneous fat that were similar to normal-weight individuals with healthy metabolomes. Individuals with a mBMI prediction that was substantially higher than their actual BMI had levels for these traits that were similar to those of obese individuals with obese metabolomes. Evaluating these data from a more clinical perspective, with individuals separated into clinical categories such as normal BMI with obese metabolome and obese BMI with healthy metabolome, generally confirmed these effects (**Figure 3** and Figure S3). Our findings suggest that the metabolome can be used as a clinically meaningful instrument, where obesity is analyzed in the context of its metabolome perturbation rather than just on BMI alone. Thus, our results are important in the frame of the current debate on the metabolically “healthy” obese and also for the identification of individuals with normal BMI^5^ but poor metabolic health^20,21^.

**Figure 3.**
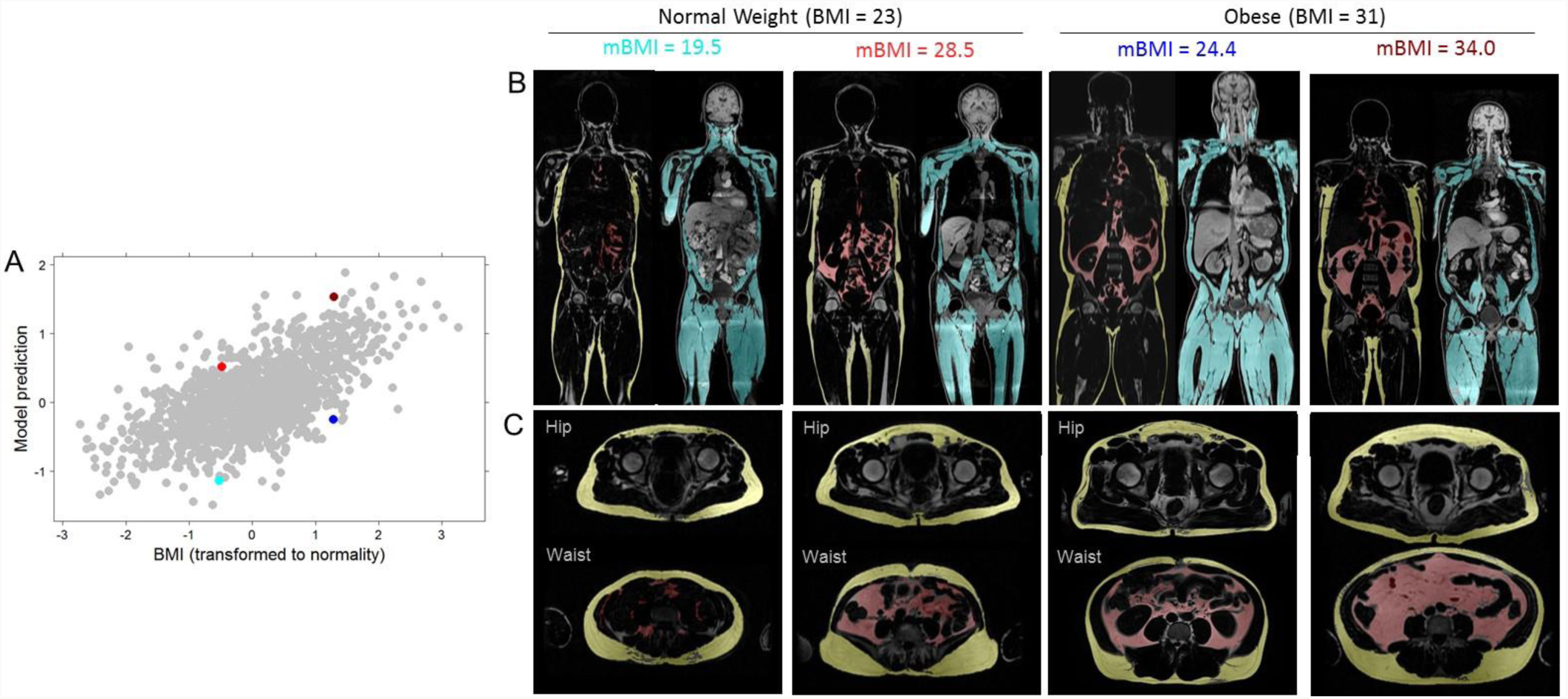
Body composition profiles from Dixon Magnetic Resonance Imaging for four outlier individuals. (A) Correlation between ridge regression model prediction of BMI and actual BMI for all unrelated individuals of European ancestry in the TwinsUK and HN dataset. Outliers highlighted in panels B and C are marked with corresponding colors. All individuals highlighted are from the outlier mBMI >> BMI or mBMI << BMI categories shown in Figure 2. (B) Body composition profiles (Red = Visceral Adipose Tissue, Yellow = Subcutaneous Adipose Tissue, Cyan = Muscle). (C) Waist to hip cross sections (Hip = Mid femoral head; Waist = Top of ASIS). (C) Identity of the individuals depicted in panels A and B.

Having characterized these outliers, we revisited their metabolome differences. As expected, those with mBMI<<BMI significantly differed in their metabolite levels from those with mBMI>>BMI for most of the 49 BMI-associated metabolites. However, two of the BMI-associated metabolites did not differ between these two groups: asparagine and cortisone. We additionally investigated the association between each of the BMI-associated metabolites and insulin resistance, as many previously reported markers of obesity have also been markers of diabetes^4,7^. We had quantitative insulin resistance measurements for 515 unrelated, European-ancestry participants. After controlling for BMI, we found that 12 of the 49 BMI-associated metabolites were also significantly associated (correcting for 49 tests requires p<0.001) with insulin resistance, all with positive directions of effect: tyrosine, alanine, kynurenate, gamma-glutamyltyrosine, 1-oleoyl-3-linoleoyl-glycerol (18:1/18:2), and six phospholipids, and as expected, glucose (see Table S1). Mannose, which recently underwent extensive study with regard to insulin resistance^14^, was nominally associated with insulin resistance after controlling for BMI in our study, p=0.004.

#### Evolution of obesity and metabolome clinical profiles

Given recent work suggesting that obese individuals who are metabolically healthy may remain at higher risk of negative health outcomes than are normal weight individuals who are metabolically healthy^20^, we next asked whether the outlier groups were more likely to become obese over time. Focusing on the 1,458 individuals from TwinsUK who had weight measurements at all three timepoints, we found that those who had a mBMI that was higher than their BMI were marginally more likely to gain weight and convert to an obese phenotype (BMI>30) over the 8-18 years of follow up. For example, 32.8% of those of normal weight but with an overweight or obese metabolome converted to being overweight or obese by timepoint 3 compared to 24.8% of those who were of normal weight and had a healthy metabolome (p=0.02, **Figure 4 and** Figure S4). Overall, the mBMI states of the individuals remained fairly stable with time and were a function of BMI changes (**Figure 4** and Figure S4). For example, 68% of the individuals who began the study with an obese metabolome ended the study with an obese metabolome. When an individuals’ weight increased and then decreased, their mBMI followed suit, and no single metabolite was significantly predictive of subsequent BMI changes (Figure S4). In summary, our results are consistent with a favorable long-term health benefit for the overweight and obese individuals with a healthy metabolome.

**Figure 4.**
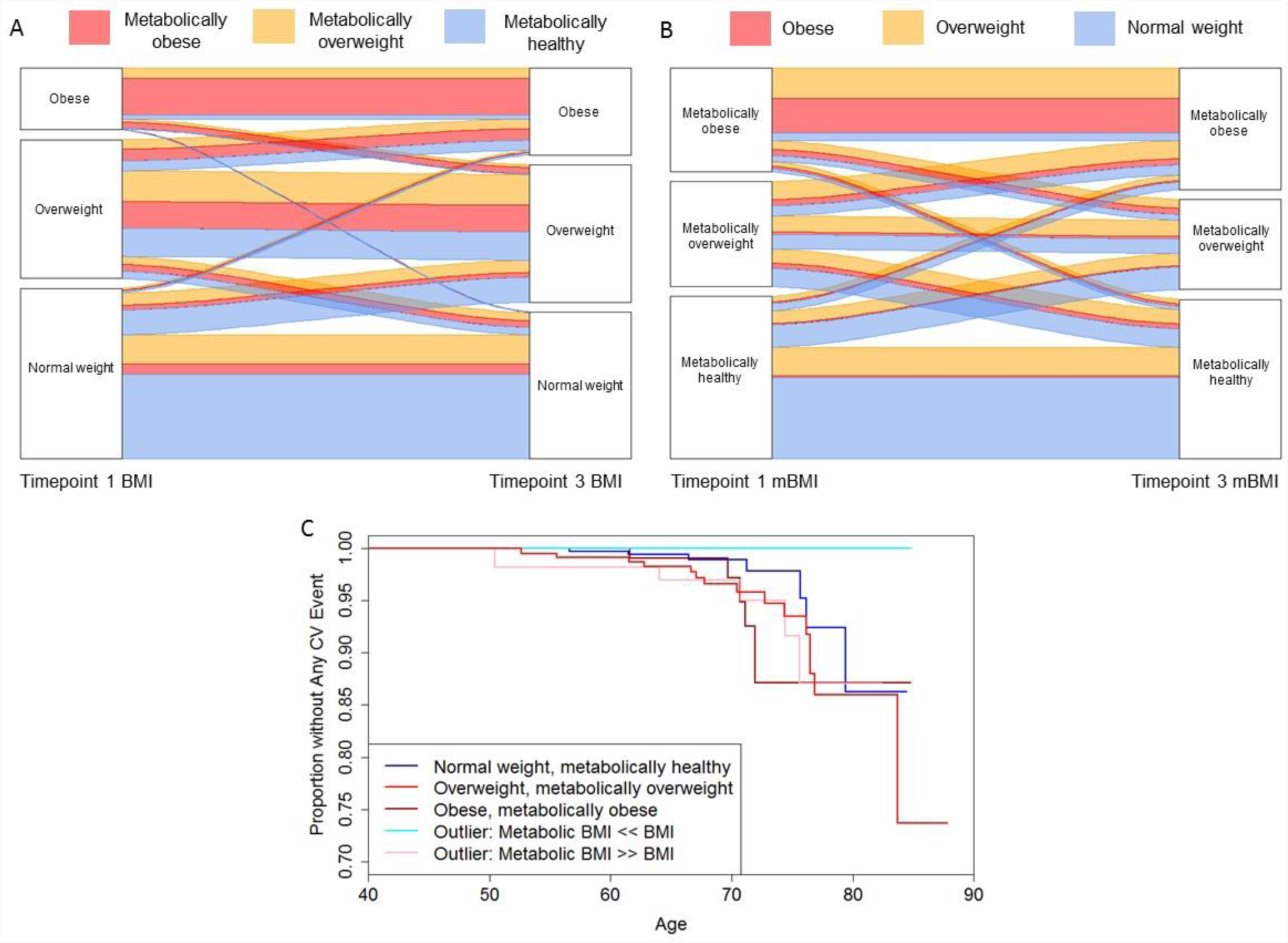
Progression of different mBMI/BMI categories. A) Alluvial plot showing the proportion of participants who remained in the same weight category or transitioned to a different weight category over the course of the 8-18 years of the TwinsUK study. Red individuals have an obese metabolome, orange individuals have an overweight metabolome, and grey individuals have a normal metabolome. B) Alluvial plot showing the proportion of participants who remained in the same mBMI category or transitioned to a different mBMI category over the course of the 8-18 years of the TwinsUK study. Red individuals begin the study with an obese BMI, orange overweight, and grey normal weight. C) Survival plot showing age until cardiac event (infarction, angina, or angioplasty). The plot is divided into those whose mBMI corresponds with their BMI (normal weight, overweight, and obese categories) as well as the two outlier groups: those with mBMI << BMI and those with mBMI >> BMI (p=0.02 for a difference between these categories in cardiovascular outcomes).

#### Cardiovascular disease outcomes

Obesity is considered a risk factor for cardiovascular disease and ischemic stroke^22^. The longitudinal nature of the TwinsUK study allowed the collection of clinical endpoints in these unselected participants. The age of participants at the first visit ranged from 33 to 74 years old (median 51); and 42 to 88 years old (median 65) at the last visit. During the follow up (median 13 years), the study recorded 53 cardiovascular events (myocardial infarct, angina, angioplasty) or strokes for 1573 individuals. We calculated that our study had 80% power to identify effects with a hazard ratio of at least 1.5 for differences in cardiovascular event outcomes between the different mBMI/BMI groups. We found that participants with a healthy metabolome (normal BMI or obese) had 2 events per hundred individuals. Individuals with an obese metabolic profile, mBMI, had 3.7 (normal BMI) and 4.2 events (in obese individuals) per hundred individuals. Separated analysis of the various endpoints confirmed the trends, more accentuated for cardiovascular than for diagnosis of stroke (Figure S5). We then performed a formal survival analysis for participants to have any cardiovascular event after the first timepoint, and we found those with healthier metabolomes to have fewer/later cardiac events (p=0.02, **Figure 4**).

#### Correlations between twins

Because twin studies are important to analyze the heritability of traits, we reassessed the BMI model predictions and obesity status of 350 sets of twins where either both twins had normal BMI (n=244), both twins were obese (n=67), or one was obese and the other had normal BMI (n=39). To keep the categories clear, individuals with BMIs between 25 and 30 (overweight) and their twins were excluded. As asserted by the model’s high specificity and sensitivity, the metabolite-based obesity predictions reflected the actual obesity status of the individuals. This was even the case when only one twin was obese: the obese twin was generally predicted by their metabolome to be obese, while the normal weight twin was not (Figure S2). The correlations between the metabolite-based obesity predictions was also substantially higher between the monozygotic twins than the dizygotic twins, as expected. Interestingly, we identified 3 sets of twins where both twins were predicted from the metabolome to be of normal weight, but both were obese, and 8 sets of twins where the reverse was true. These outliers were thought to represent healthy obese and normal weight, metabolically unhealthy individuals described above.

### Genetic analyses

#### Known genetics of obesity

We first investigated the known genetic factors contributing to high BMI. We calculated polygenic risk scores for BMI using known associations from the considerable literature of obesity and BMI GWAS^23^. As previously reported, we found that polygenic risk score only explained 2.2% of the variation in BMI at each of the three TwinsUK timepoints and in Health Nucleus for unrelated participants of European ancestry (Figure S6). We investigated whether unique individuals with the highest polygenic risk scores would have a significant perturbation of the metabolome and anthropomorphic, insulin resistance and DEXA measurements (**Figure 5**). While the data did not support a strong role for polygenic risk, there were trends for higher polygenic risk scores to be associated with a higher android/gynoid ratio (p=0.04) and waist/hip ratio (p=0.04). However, there was no statistical association between the polygenic score and mBMI (p=0.16).

**Figure 5.**
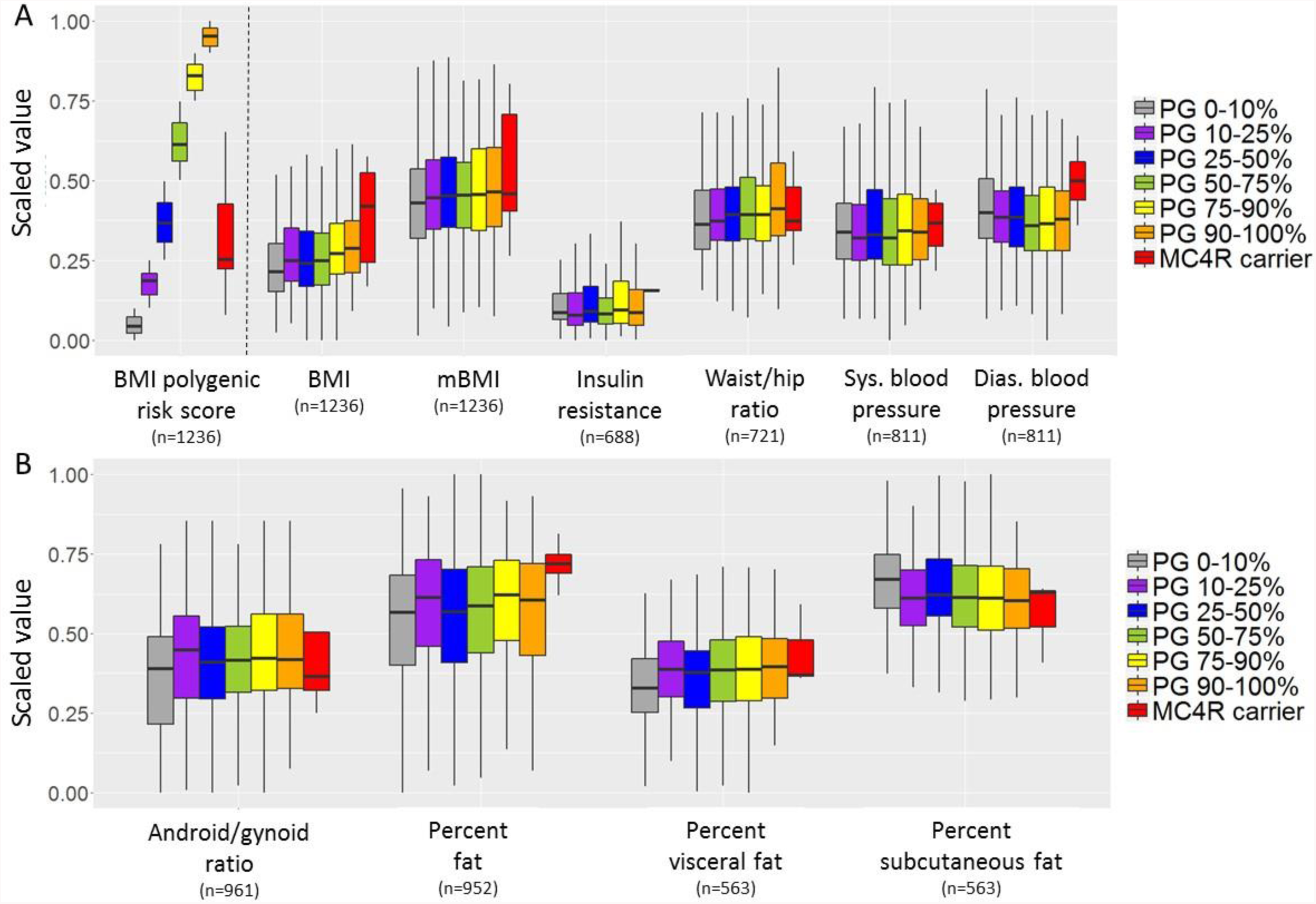
Genetic risk compared to BMI-relevant variables. (A) Correlation between polygenic risk score (PG) category, *MC4R* carrier status, and BMI and anthropomorphic and clinical measurements for all unrelated individuals of European ancestry in the TwinsUK and HN dataset. All y-axis values are scaled to a range from 0-1 to allow comparison across groups. The same process is used in (B) to show DEXA imaging values. While there was a trend for genetic risk to be associated with various measurements, the polygenic risk score achieved nominal p<0.05 for BMI, waist/hip ratio and android/gynoid ratio, and *MC4R* carrier status achieved nominal p<0.05 for BMI.

Studies of rare variants in obesity have identified *MC4R* mutations as having effects large and clear enough to be appropriate for study in our dataset^24^. We therefore identified members of the study populations who were carrying rare (MAF<0.01%) coding variants in the known obesity gene melanocortin 4 receptor (*MC4R*). We identified 8 such carriers in the subset of unrelated participants (**Table 2**). Each variant was observed in one unrelated individual, and 5 of the 8 had already been annotated as causing obesity in clinical databases HGMD or ClinVar (**Table 2**). As a group, *MC4R* carriers had significantly higher BMI (p=0.02) than did non-carriers as well as non-significant trends toward a higher diastolic blood pressure, insulin resistance, and percent body fat (**Figure 5**). However, not all rare variants may be deleterious, and the metabolic impact could have been greater for the true subset of functional variants. The BMI data in the participants supported a pathogenic role for five of the variants (Met292fs, Arg236Cys, Ser180Pro, Ala175T, and Thr11Ala), but did not corroborate a role of Ile170V, which is defined in HGMD and ClinVar as pathogenic^25,26^. Importantly, of the five sets of twins who both carried the same *MC4R* variant, three sets included twins who were both overweight or obese. In the two cases where a carrier’s twin did not have the *MC4R* variant, their BMI was lower than their twin’s. We observed an enrichment of *MC4R* variant carriers among obese individuals with low polygenic risk scores (Figure S6). Out of 31 participants who were obese with polygenic risk scores in the lowest quartile, 6.1% were *MC4R* variant carriers, while the carrier frequency was just 0.3% in those of normal weight.

**Table 2.** Variants identified in *MC4R* in unrelated participants of European ancestry.

| Variant | Protein change | Study MAF | Global gnomad MAF | Known obesity annotation | Carrier BMI | Carrier twin BMI | Twin non-carrier BMI | Twin zygosity |
| --- | --- | --- | --- | --- | --- | --- | --- | --- |
| chr18:60371541 G/A | p.Ser270Phe | 0.036% | 0.003% | None | 25.7 | 24.8 | N/A | MZ |
| chr18:60372307 G/A | p.Leu15Phe | 0.036% | 0.000% | None | 23 | 22.6 | N/A | MZ |
| chr18:60371474 CA/C | p.Met292fs | 0.036% | <0.003% | None | 32.8 | N/A | 28.8 | DZ |
| chr18:60371644 G/A | p.Arg236Cys | 0.036% | 0.003% | HGMD highC DM | 34.5 | 34.5 | N/A | DZ |
| chr18:60371812 A/G | p.Ser180Pro | 0.036% | <0.003% | ClinVar LP | 34.2 | 34.4 | N/A | DZ |
| chr18:60371827 C/T | p.Ala175Thr | 0.036% | 0.019% | ClinVar P and HGMD highC DM | 29 | 28.5 | N/A | MZ |
| chr18:60371842 T/C | p.Ile170Val | 0.036% | 0.013% | ClinVar P and HGMD highC DM | 22.6 | N/A | 21.3 | DZ |
| chr18:60372319 T/C | p.Thr11Ala | 0.036% | <0.003% | HGMD lowC DM | 36 | N/A | N/A | N/A |

MAF = minor allele frequency; HGMD highC DM = Human Gene Mutation Database high-confidence disease-causing mutation; lowC = low-confidence; LP = likely pathogenic; P = pathogenic; MZ = monozygotic; DZ = dizygotic. Each variant was only seen once in the unrelated participants of this study.

#### Genetics of the metabolically healthy obese

We obtained additional support of the decoupling of the genetics of high BMI versus the basis of obesity and predicted mBMI from the analysis of outliers. Individuals with a mBMI that was substantially lower than their actual BMI had a higher polygenic risk score for BMI than did other groups. In contrast, those whose mBMI was substantially higher than their actual BMI had low polygenic risk scores (**Figure 2B**; p=0.006 for a difference between these two groups). This result would support the notion that the polygenic risk score for BMI may capture an anthropomorphic phenotype (larger-framed individuals) rather than a unique association with obesity as a disease trait.

#### Genetics of metabolome differences

Last, we investigated whether obese individuals with different genetic backgrounds had different metabolomes from other obese individuals. We first searched for metabolites that could distinguish individuals with different BMI polygenic risk scores or *MC4R* variant carriers. Linear regression showed no significant associations between any single metabolites and polygenic risk or *MC4R* carrier status in either the entire population or in only the obese individuals. This result implies that metabolites are unlikely to be intermediate phenotypes that explain the underlying genetics of obesity. To check for more specific signals beyond the compiled polygenic risk score, we also performed separate analyses of each of the 97 variants that are used to calculate the polygenic risk score. We found no evidence for any of these known GWAS variants to be more strongly associated with a metabolite than with BMI itself, though our power for discovery was limited given the very small effect sizes of most individual GWAS variants. In summary, although it is known that there is a strong genetic component to metabolite levels^27^, most of the metabolic perturbations that occur in the obese state are a response to obesity as opposed to shared genetic mechanisms.

## Discussion

The results of the present study highlight the profound disruption of the metabolome in obesity and identifies a metabolome signature that serves to examine metabolic health beyond anthropomorphic measurements. Nearly one third of the approximately 1000 metabolites measured in the study were associated with BMI, and 49 were selected as a strong signature for the study of the relationship between BMI, obesity, metabolic disease and the genetics of BMI.

Consistent with previous studies and earlier work in the TwinsUK cohort, branched-chain and aromatic amino acids, and metabolites involved in nucleotide metabolism, such as urate and pseudouridine, are strongly perturbed by obesity^4,6,7,9^. The underlying reason for the perturbation of branched-chain amino acid metabolism in obese individuals and those with insulin resistance is thought to be related to differences in the amino acid catabolism in adipose tissue^28^. The single metabolite with the most significant association with BMI was urate, as we previously reported^9^. It is well known that uric acid increases with obesity, due to insulin resistance reducing the kidneys’ ability to eliminate uric acid, but previous work has not emphasized the power of urate to predict BMI^6,7,29^. We also found a strong signal for lipids to be associated with BMI, with an enrichment of associations found for glycerol lipids. These results are consistent with previous studies showing that sphingomyelins and diacylglycerols increase with BMI while lysophosphocholines decrease with BMI, with other various phosphatidylcholines having effects in both directions^6,7,4^. A number of BMI-associated metabolites (12 of the 49-metabolite signature) were associated with insulin resistance after controlling for BMI. As previously observed^20,30–32^, the metabolome abnormalities associated with high BMI corrected with loss of weight. However, our study found that metabolite levels did not provide predictive power for future weight changes (Figure S4). Overall, the metabolome perturbations appear as a consequence of changes in weight as opposed to being a contributing factor.

The metabolome signature identified individuals whose predicted mBMI was either substantially lower or higher than their actual BMI. These individuals include the metabolically healthy obese, but we also emphasize the importance of the metabolome anomalies in identifying unhealthy individuals with a normal BMI. These profiles were generally stable over the prolonged follow-up. An abnormal metabolome signature, irrespective of BMI, was associated in the present study with three-fold increase in cardiovascular events (**Figure 6**). Thus, while our findings are in line with the known relationships between metabolically healthy obese status and health-related traits like metabolic syndrome and body fat^20,33,34^, we extend this relationship to the broader category of metabolically healthy and unhealthy individuals on the basis of the disparity between mBMI and BMI. For example, we observed differences in waist/hip ratio, percent visceral fat, and blood pressure between mBMI/BMI outliers despite having the same BMI distribution. The fact that the metabolically healthy obese have a high BMI polygenic risk score also supports the concept that some of the genetic studies may capture anthropomorphic associations – body size - rather than obesity *sensu stricto*. Overall, the health consequences observed across the various mBMI groups indicate that there is a durable benefit of maintaining a healthy metabolome signature and points to an ongoing risk for the individuals that have an unhealthy metabolome despite stability of BMI.

**Figure 6.**
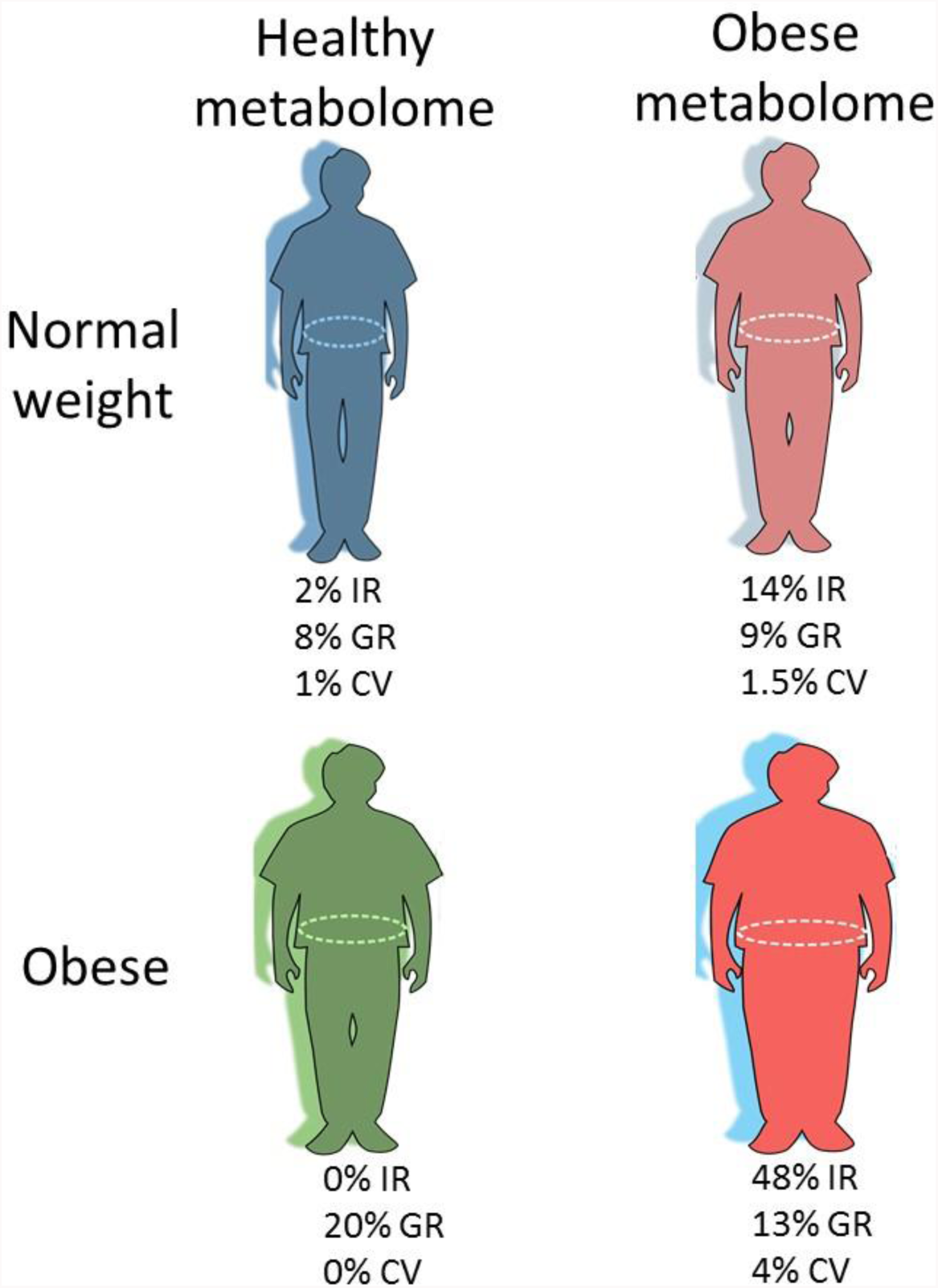
Representative clinical phenotypes of mBMI/BMI outliers. While there is a continuum of obesity and metabolic perturbations, there are four representative extant phenotypes that are schematically represented in the figure. Indicated are salient features of these groups: rates of insulin resistance (IR) at timepoint 1, high BMI genetic risk (GR, top decile of polygenic risk or MC4R carrier), and rates of cardiovascular events (CV) during the study follow up.

In contrast with metabolomics analyses, the present study does not support a strong association between metabolome changes and the genetics of BMI defined by a 97-variant polygenic risk score^23^. This may be explained by the fact that known BMI GWAS loci explain only a small fraction (∼3%) of BMI heritability^23^. Despite this overall lack of explanatory power, there was a clear signal for individuals with higher polygenic risk scores to have greater rates of obesity. Because the genetic risk score does not include rare variants, we also identified individuals who carried rare functional variants in the known obesity gene *MC4R*, which is the single best example of a gene where rare coding variants have a large effect on obesity^24^. The carriers of these variants were often obese individuals, but their metabolome was not categorically different from that of other obese individuals. The lack of metabolome differences for carriers of variants in this gene is not surprising given that *MC4R* variants cause obesity by increasing appetite. However, we did find that obese carriers of *MC4R* variants often had low polygenic risk scores for obesity; out of 31 participants who were obese with polygenic risk scores in the lowest quartile, 6.1% were *MC4R* variant carriers, while the carrier frequency was just 0.3% in those of normal weight. Thus, our study shows the interest of sequencing obese individuals with low polygenic risk scores because of the apparent enrichment for monogenic contributions. As we completed this study, a large consortium provided additional detail on the role of variants in pathways that implicate energy intake and expenditure in obesity^24^.

In summary, the present study highlights the health risks of the perturbed metabolome. The study also decouples the genetics of BMI from metabolic health and serves to prioritize a subset of individuals for genetic analysis. The assessment of the metabolome and genome of BMI lays groundwork for future studies of the heterogeneity of obesity and treatment of its endophenotypes. Specifically, the metabolome signature can act as a biomarker of response to the new therapeutics that target patients with *MC4R* mutations^35^. In the future, metabolic profiling could help select patients for clinical trials beyond genetic sequencing, thus expanding drug utility^11^.

### STAR+ Methods

#### Samples and study design

Our study included 1,969 European ancestry twins enrolled in the TwinsUK registry, a British national register of adult twins^12^. We previously reported a detailed study of the genetic variants influencing the human metabolome in this cohort^27^. Serum samples were collected at three visits, 8-18 (median 13) years apart. The cohort is mainly composed of females (96.7%), and the sample set we used included 388 monozygotic twin pairs, 519 dizygotic twin pairs, and 155 unrelated individuals. The age of participants at the first timepoint ranged from 33 to 74 years old (median 51); 36 to 81 years old (median 59) at the second timepoint; and 42 to 88 years old (median 65) at the third timepoint. The BMI values measured at each metabolome timepoint were taken within two years of the blood draw date. The twins study was approved by St. Thomas’ Hospital Research Ethics Committee, and all participants provided informed written consent. BMI data were available for 1743 participants within two years of the timepoint for metabolome timepoint 1, 1834 for within two years of timepoint 2, and 1777 for up to 2 years before timepoint 3 or 4 years after this timepoint; 1,458 individuals had all three datapoints.

For independent validation and studies of phenotypes correlated with metabolic BMI outliers, we enrolled 617 unselected adults more than 18 years old who were able to come to the Health Nucleus in La Jolla, CA for a clinical research protocol^13^. Participants underwent a verbal review of the institutional review board-approved consent (Western Institutional Review Board). Participants ranged in age from 18-89 years old (median 53), were 32.9% female, and had BMI data measured at one timepoint.

#### Phenotyping

Individuals in the TwinsUK cohort and Health Nucleus both underwent DEXA imaging. The data from these scans were used to calculate android/gynoid ratio, percent body fat, visceral fat, and subcutaneous fat. TwinsUK cohort participants were additionally measured for circumference at the waist and hip using a measuring tip to calculate the waist/hip ratio. For a selected number of Health Nucleus participants, images of fat and water (imaging of muscle) were available from symmetrical chemical shift Magnetic Resonance Imaging (MRI) via the Dixon method. Quantitative insulin resistance (homeostatic model assessment, HOMA) was calculated as fasting insulin x fasting glucose / 405, and being insulin resistant was defined by HOMA score ≥3 (http://gihep.com/calculators/other/homa/)^36^.

#### Metabolite Profiling

The non-targeted metabolomics analysis of 901 metabolites in the TwinsUK cohort and 1,007 metabolites in the Health Nucleus cohort was performed at Metabolon, Inc. (Durham, North Carolina, USA) on a platform consisting of four independent ultra high performance liquid chromatography-tandem mass spectrometry (UPLC-MS/MS) methods. The detailed descriptions of the platform can be found in our previous publications^27,37^. For the TwinsUK cohort, blood serum was used for analysis, and the resulting raw values were transformed to z scores using the mean and standard deviation. For the Health Nucleus cohort, blood plasma was used for analysis, and values from multiple experimental batches were normalized into Z-scores based on a reference cohort of either 42 (n=457) or 300 (n=176) self-reported healthy individuals run with each batch. The 42 and 300-normalized batches were converted to the same scale using linear transformation based on the values obtained from 7 runs that included both the 42 and 300 controls. Samples with metabolite measurements that were below the detection threshold were imputed as the minimum value for that metabolite.

#### Genome sequencing and analysis

As previously described^38^, DNA samples were sequenced on an Illumina HiSeqX sequencer utilizing a 150 base paired-end single index read format. Reads were mapped to the human reference sequence build HG38. Variants were called using ISIS Analysis Software (v. 2.5.26.13; Illumina). A linear mixed model was applied to account for family structure in the cohort while testing for associations between genetic variants and the different phenotypes: BMI; BMI prediction model values and residuals after accounting for BMI, age, sex; and levels of the 49 BMI-associated metabolites. A genetic similarity matrix (GSM) was constructed from 301,556 variants that represented a random 20% of all common (MAF>5%) variants genome-wide after linkage-disequilibrium (LD) pruning (r^2^ less than 0.6, window size 200 kb) and was used to model the random effect in the linear mixed model via a “leave-out-one-chromosome” method for each tested variant. Each of 97 known BMI-associated variants was tested independently using customized Python scripts wrapping the FaST-LMM package^39,23^. Principal component axes were calculated to check ethnicity using plink, and the first principal component for those of European ancestry was used as a covariate in analyses of unrelated individuals in R described below. Polygenic risk scores were calculated using genotypes for 97 variants whose associations and betas had been published previously^23^. Rare variants in the gene *MC4R* were defined as coding and splice variants with MAF<0.1%.

#### Statistical analysis

R was used for the analysis and data manipulation. Bonferroni correction was used for all analyses. For each quantitative analysis of BMI or other traits, the subset of BMI values or other outcome variables used were rank-ordered and forced to a normal distribution. Analyses comparing metabolites to BMI were performed in R using the lm function, and age, sex, and the first genetic principal component were included as covariates. The obesity prediction model was built using ridge regression (alpha=0) with glmnet in R. The residuals used to separate participants into the five categories shown in Figure 2 were calculated using age, sex, and initial BMI. Heatmaps were generated in R using the pheatmap package. Survival analysis was performed using coxph in R with age at first visit included as a covariate. Power calculation was performed using the power.stratify command in powerSurvEpi.

## Acknowledgements

We thank E. Muse for valuable comments. We acknowledge N. Schenker-Ahmed, L. Huang, M. Tyagi, and P. Sheth for aiding with data collection. Twins UK receives funding from the Wellcome Trust. European Community’s Seventh Framework Programme (FP7/2007-2013 and Horizon 2020 to TwinsUK); the National Institute for Health Research (NIHR) Clinical Research Facility at Guy’s & St Thomas’ NHS Foundation Trust and NIHR Biomedical Research Centre based at Guy’s and St Thomas’ NHS Foundation Trust and King’s College London.

## Author contributions statement

A.T. and E.C. conceived of the experiment(s), E.C. conducted the experiment(s), L.G. supported metabolome analyses and interpretation, E.C., J.C.V. and A.T. analyzed the results. C.T.C. and B.T. provided detailed insight into metabolic diseases, C.S., N.S, L.H., E.K. and L.N. aided with data collection and interpretation, T.D.S. is responsible for the TwinUK study. All authors reviewed the manuscript.

## Competing financial interests

E.C., C.S., N.S, L.H., E.K. L.N. and J.C.V are employees of Human Longevity, Inc. L.G. is employee of Metabolome Inc. A.T., B.T., and T.S. declare no conflict of interest.

## References

1. MacMahon, S. et al. Body-mass index and cause-specific mortality in 900 000 adults: Collaborative analyses of 57 prospective studies. Lancet 373, 1083–1096 (2009).

2. Hales, C. M., Carroll, M. D., Fryar, C. D. & Ogden, C. L. Prevalence of Obesity Among Adults and Youth: United States, 2015-2016. NCHS Data Brief (2017).

3. WHO. Obesity and overweight. Fact sheet (2017). Available at: http://www.who.int/mediacentre/factsheets/fs311/en/.

4. Park, S., Sadanala, K. C. & Kim, E.-K. A Metabolomic Approach to Understanding the Metabolic Link between Obesity and Diabetes. Mol. Cells 38, 587–596 (2015).

5. Chen, H. H. et al. The metabolome profiling and pathway analysis in metabolic healthy and abnormal obesity. Int. J. Obes. 39, 1241–1248 (2015).

6. Butte, N. F. et al. Global metabolomic profiling targeting childhood obesity in the Hispanic population. Am. J. Clin. Nutr. 102, 256–267 (2015).

7. Ho, J. E. et al. Metabolomic Profiles of Body Mass Index in the Framingham Heart Study Reveal Distinct Cardiometabolic Phenotypes. PLoS One 11, e0148361 (2016).

8. Piening, B. D. et al. Integrative Personal Omics Profiles during Periods of Weight Gain and Loss. Cell Syst. 1–14 (2018). doi:10.1016/j.cels.2017.12.013

9. Menni, C. et al. Metabolomic Profiling of Long-Term Weight Change: Role of Oxidative Stress and Urate Levels in Weight Gain. Obesity (Silver Spring). 25, 1618–1624 (2017).

10. Neeland, I. J., Poirier, P. & Després, J.-P. Cardiovascular and Metabolic Heterogeneity of Obesity. Circulation 137, 1391–1406 (2018).

11. Yanovski, S. Z. & Yanovski, J. A. Toward Precision Approaches for the Prevention and Treatment of Obesity. JAMA 319, 223–224 (2018).

12. Moayyeri, A., Hammond, C. J., Hart, D. J. & Spector, T. D. The UK Adult Twin Registry (TwinsUK Resource). Twin Res. Hum. Genet. 16, 144–149 (2013).

13. Perkins, B. A. et al. Precision Medicine Screening Using Whole Genome Sequencing And Advanced Imaging To Identify Disease Risk In Adults. bioRxiv 133538 (2017). doi:10.1101/133538

14. Lee, S. et al. Integrated Network Analysis Reveals an Association between Plasma Mannose Levels and Insulin Resistance. Cell Metab. 24, 172–184 (2016).

15. Wikoff, W. R. et al. Metabolomics analysis reveals large effects of gut microflora on mammalian blood metabolites. Proc. Natl. Acad. Sci. 106, 3698–3703 (2009).

16. Björntorp, P. & Rosmond, R. Obesity and cortisol. in Nutrition 16, 924–936 (2000).

17. Praveen, E. P. et al. Morning cortisol is lower in obese individuals with normal glucose tolerance. Diabetes, Metab. Syndr. Obes. Targets Ther. 4, 347 (2011).

18. Walker, B. R., Soderberg, S., Lindahl, B. & Olsson, T. Independent effects of obesity and cortisol in predicting cardiovascular risk factors in men and women. J Intern Med 247, 198–204 (2000).

19. Wirix, A. J. G. et al. Is there an association between cortisol and hypertension in overweight or obese children? JCRPE J. Clin. Res. Pediatr. Endocrinol. 9, 344–349 (2017).

20. Caleyachetty, R. et al. Metabolically Healthy Obese and Incident Cardiovascular Disease Events Among 3.5 Million Men and Women. J. Am. Coll. Cardiol. 70, 1429–1437 (2017).

21. Molli, A. E. I. et al. Metabolically healthy obese individuals present similar chronic inflammation level but less insulin-resistance than obese individuals with metabolic syndrome. PLoS One 12, (2017).

22. Rhee, E. Being Metabolically Healthy, the Most Responsible Factor for Vascular Health. Diabetes Metab J 42, 19–25 (2018).

23. Locke, A., Kahali, B., Berndt, S., Justice, A. & Pers, T. Genetic studies of body mass index yield new insights for obesity biology. Nature 518, 197–206 (2015).

24. Turcot, V. et al. Protein-altering variants associated with body mass index implicate pathways that control energy intake and expenditure in obesity. Nat. Genet. 50, 26–35 (2018).

25. Landrum, M. J. et al. ClinVar: Improving access to variant interpretations and supporting evidence. Nucleic Acids Res. 46, D1062–D1067 (2018).

26. Stenson, P. D. et al. Human Gene Mutation Database (HGMD??): 2003 Update. Human Mutation 21, 577–581 (2003).

27. Long, T. et al. Whole-genome sequencing identifies common-to-rare variants associated with human blood metabolites. Nat. Genet. 49, 568–578 (2017).

28. Newgard, C. B. Interplay between lipids and branched-chain amino acids in development of insulin resistance. Cell Metab. 15, 606–614 (2012).

29. Facchini, F., Chen, Y. D., Hollenbeck, C. B. & Reaven, G. M. Relationship between resistance to insulin-mediated glucose uptake, urinary uric acid clearance, and plasma uric acid concentration. Jama 266, 3008–3011 (1991).

30. Oberbach, A. et al. Combined proteomic and metabolomic profiling of serum reveals association of the complement system with obesity and identifies novel markers of body fat mass changes. J. Proteome Res. 10, 4769–4788 (2011).

31. Reinehr, T. et al. Changes in the serum metabolite profile in obese children with weight loss. Eur. J. Nutr. 54, 173–181 (2014).

32. Wahl, S. et al. Multi-omic signature of body weight change: results from a population-based cohort study. BMC Med. 13, 48 (2015).

33. Karelis, A. D. et al. The metabolically healthy but obese individual presents a favorable inflammation profile. J. Clin. Endocrinol. Metab. 90, 4145–4150 (2005).

34. Brochu, M. et al. What are the physical characteristics associated with a normal metabolic profile despite a high level of obesity in postmenopausal women? J. Clin. Endocrinol. Metab. 86, 1020–1025 (2001).

35. Kühnen, P. et al. Proopiomelanocortin Deficiency Treated with a Melanocortin-4 Receptor Agonist. N. Engl. J. Med. 375, 240–246 (2016).

36. Matthews, D. R. et al. Homeostasis model assessment: insulin resistance and beta-cell function from fasting plasma glucose and insulin concentrations in man. Diabetologia 28, 412–419 (1985).

37. Cohen, I. V. et al. Acetaminophen (Paracetamol) Use Modifies the Sulfation of Sex Hormones. EBioMedicine 0, 1–8 (2018).

38. Telenti, A. et al. Deep sequencing of 10,000 human genomes. Proc. Natl. Acad. Sci. 113, 11901–11906 (2016).

39. Lippert, C. et al. FaST linear mixed models for genome-wide association studies. Nat. Methods 8, 833–835 (2011).

